# Human Endogenous Retrovirus Expression is Dynamically Regulated in Parkinson’s Disease

**DOI:** 10.1101/2023.11.03.565438

**Authors:** Juozas Gordevičius, Thomas Goralski, Alexis Bergsma, Andrea Parham, Emily Kuhn, Lindsay Meyerdirk, Mitch McDonald, Milda Milčiūtė, Elizabeth Van Putten, Lee Marshall, Patrik Brundin, Lena Brundin, Viviane Labrie, Michael Henderson, John Andrew Pospisilik

## Abstract

Parkinson’s disease (PD) is a progressive, debilitating neurodegenerative disease that afflicts approximately every 1000th individual. Recently, activation of genomic transposable elements (TE) has been suggested as a potential driver of PD onset. However, it is unclear where, when, and to what extent TEs are dysregulated in PD. Here, we performed a multi-tissue transcriptional analysis of multiple patient cohorts and identified TE transcriptional activation as a hallmark of PD. We find that PD patients exhibit up-regulation primarily of human endogenous retrovirus (HERV) transcripts in prefrontal cortex tissue, prefrontal neurons as well as in blood, and we demonstrate that TE activation in the blood is highest at the time of PD diagnosis. Supporting a potentially causal association between ERV dysregulation and PD heterogeneity, reduced gene dosage of the TE repressor Trim28 triggers transcriptional changes highly correlated to those measured in animal models of synucleinopathy (PFF-injection), and importantly, to those exhibited by patients themselves. These data identify ERV up-regulation as a common feature of central and peripheral PD etiology, and highlight potential roles for Trim28-dependent TEs in stratifying and monitoring PD and treatment compliance.

## INTRODUCTION

Parkinson’s disease (PD) is the most common neurodegenerative movement disorder, with an incidence rate of 108 to 212 per 100,000 among persons ages 65 and older^1^. While the molecular etiology of PD is not well understood, PD is characterized by progressive degeneration of the dopaminergic neurons of the substantia nigra and the appearance of intraneuronal inclusions enriched in misfolded α-synuclein (Lewy bodies and Lewy neurites) ^2^. The pathogenesis of PD includes typically a long prodromal phase (lasting several years) before onset of hallmark symptoms, and evidence indicates that the disease is rooted in both genetic and environmental factors^3^. Familial PD is associated with strong disease predisposing mutations and accounts for 3-5% of all cases^4^. The underpinnings of sporadic PD, on the other hand, have been more elusive. Estimates from twin studies indicate an ∼20% heritability^5^; genome-wide association studies (GWAS) similarly place it at ∼16-36%^6^. What is clearly acknowledged in the field is that substantial phenotypic heterogeneity is observed both for familial forms (e.g. including individuals with mutation who do not get disease) and for idiopathic forms (e.g. many PD patients fail to exhibit substantial known polygenic risk and vice versa). These data suggest a strong role for environmental and epigenetic mechanisms in PD etiology.

Recently, transposable elements (TEs) were postulated to play a role in PD^7^. TEs are mobile DNA elements which originate from viruses that integrated into our ancestral germline DNA millions of years ago^8^. TEs have the ability to replicate and re-insert, causing de novo insertional mutations. TE presence in all plant and animal genomes is believed to highlight their critical role in driving evolution, speciation and the emergence of novel trait regulation. Specifically, TEs can contribute novel cis-regulatory elements (e.g., enhancers, alternate promoters) and provide a multi-copy sequence basis for transcriptional co-regulation^9^. While TEs are mostly epigenetically silenced, activation of retrotransposons such as endogenous retroviruses (ERVs), a subclass of TEs, can result in genomic and cellular instability, disruption of coding regions, and modulation of epigenetic and post-transcriptional regulation of gene expression^9^. Loss of silencing at TEs represents a mechanism to release cryptic genetic information and has been associated with activation of viral immune response pathways within cells^10,11^. A recent publication demonstrated an increase of retrotransposition events in PD blood, suggesting that retrotransposons may be associated with PD risk and disease progression^12^.

Human ERVs (HERVs) represent ∼8% of the human genome. They originate from the cumulative infection and integration of germline DNA by retroviruses^13^ and are kept silent by dedicated epigenetic machinery including the Trim28 silencing complex. Trim28 is considered a master regulator of ERV silencing^14,15^. A large multi-domain protein, Trim28 is recruited to genomic ERVs via Krüppel-associated box (KRAB)-zinc finger transcription factors and coordinates the assembly of a multiprotein complex to establish heterochromatin deposition through DNA methylation and the repressive histone mark H3K9me3^16^. Homozygous Trim28 deletion in mice is early embryonic lethal and conditional maternal deletion mutants exhibit highly variable developmental abnormalities^17,18^. Our group recently characterized a Trim28 haploinsufficiency that survives well into adulthood and old age and found that animals exhibit exaggerated phenotypic variation, implicating Trim28, and by extension ERV dysregulation, in the control of non-genetic non-environmental phenotypic outcomes, also known as unexpected phenotypic variation (UPV)^19^. Trim28 is highly expressed in the brain and is known to silence ERV expression in mouse and human neural progenitor cells^20,21^.

Here, we identify transcriptional activation of TEs as a hallmark of PD and show that Trim28 haploinsufficiency triggers a PD-mimicking transcriptional state in the CNS. Supporting a potentially causal association between ERV dysregulation and PD heterogeneity, reduced gene dosage of the TE silencer Trim28 triggers transcriptional changes highly correlated to those measured in animal models of synucleinopathy (intrastriatal injection of preformed fibrils of alpha-synuclein), and importantly, to those exhibited by patients themselves. We find that PD patients exhibit up-regulated ERV transcripts in prefrontal cortex (PFC) tissue, in PFC neurons, as well as in blood, using multiple cohorts. By comparing samples ipsi- and contra-lateral to symptom onset, we demonstrate that ERV up-regulation is coupled to PD severity, findings that are also consistent with the idea that neuroinflammation contributes towards PD etiology. These data identify ERV dysregulation and Trim28 haploinsufficiency as novel features of PD heterogeneity and experimental synucleinopathy. They highlight potential utility of Trim28-dependent TEs in PD diagnosis, stratification and treatment compliance.

## RESULTS

### TEs are up-regulated in PD prefrontal cortex

We performed RNA-seq of bulk prefrontal cortex (PFC) tissue from patients with PD (N = 24) and a set of healthy controls (N = 13) of Caucasian race 75.51 (± 5.20 SD) years old. Thirteen of the PD samples (termed ‘Matched’) were isolated from the brain hemisphere contralateral symptom onset. For the remaining 11 PD samples (termed ‘Unmatched’) samples were taken from the ipsilateral hemisphere relative to symptom onset. ‘Control’ samples were obtained from persons without known neurodegenerative disease. After QC and data filtering we reliably quantified the expression of 19,883 genes and 202 TEs across all the samples. After normalization and variance correction we applied a robust linear regression model to each gene and TE with and adjusted the model for age, sex, brain hemisphere, post-mortem interval, estimated excitatory neuron proportion and other possible sources of unknown variation using RUV (Risso 2014).

Focusing first on Matched PD samples, we identified 6,147 differentially expressed genes (DEGs) and 86 differentially expressed TE’s (DETEs) (**Additional File 2**), relative to controls. Interestingly, whereas DEGs were more likely to be down-regulated relative to controls, DETEs were preferentially up-regulated, suggesting a global impairment in TE silencing (**Fig. 1a**; p = 3.6 × 10^−8^ and p = 0.0085 for DEGs and DETEs, respectively). These differences were validated in a parallel analysis of ‘Unmatched’ PD samples. Interestingly, whereas DEGs showed no directional skew, TEs were once again up-regulated (OR = 4.00, p = 0.05) relative to controls (**Additional File 1: Figure S1a;** OR = 1.03; Fisher’s exact test). Consistent with the spatial progression of PD pathogenesis, the ‘Unmatched’ PD samples showed fewer DEGs (3,393 genes) and DETEs (34 TEs). These data identified TE up-regulation as a feature of both early and late PD pathology. Thus, transposable elements are derepressed in PD PFC.

**Fig. 1.**
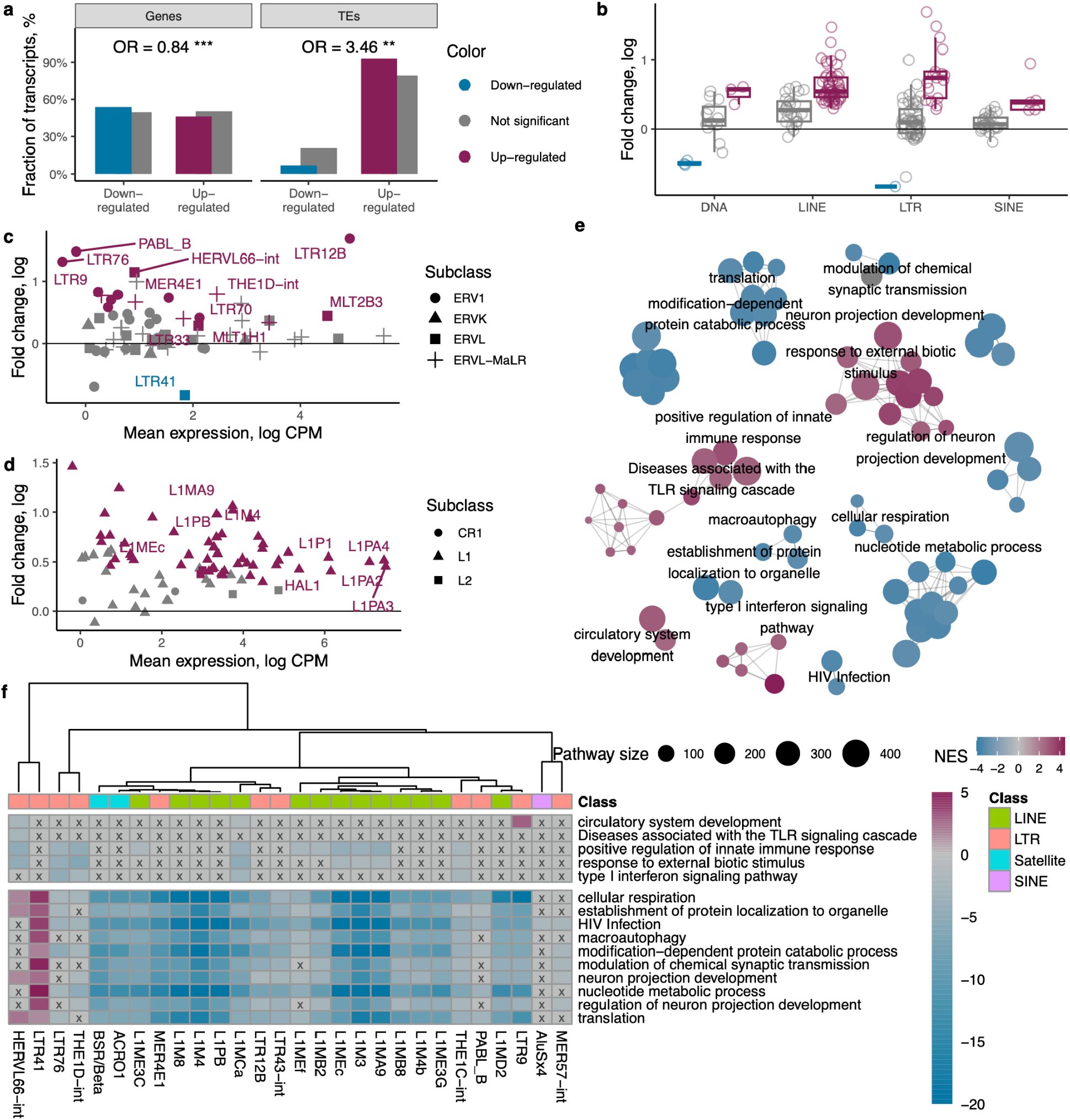
Differential expression of genes and up-regulation of transposable elements (TEs) in the bulk prefrontal cortex neuron tissue comparing Matched PD to control brain samples. **(a)** Fraction of up-regulated and down-regulated significant genes and TEs (colored fill) compared to the fraction of non-significant up-regulated and down-regulated genes as well as TEs (gray fill). **(b)** Fold change of TEs stratified by class and colored by statistical significance. **(c)** Mean expression and fold change of long terminal repeat elements (LTRs) colored by statistical significance and shaped by element subclass. **(d)** Mean expression and fold change of LINE elements colored by statistical significance and shaped by element subclass. **(e)** Network and clustering of the top 50 most up-regulated and 50 most down-regulated pathways obtained through gene set enrichment analysis. Each cluster is named after the most prominent pathway decided by the PageRank algorithm. **(f)** Mixed effects modeling of the association between the expression of TEs and genes belonging to the leading pathways indicated in clustering analysis. Fill color indicates signed log p value. P values were adjusted for multiple testing using FDR. Heatmap cells with a value exceeding FDR q-value higher than 0.05 were marked with an x and gray color.

Among the significantly affected TEs in the Matched PD group, we identified six DNA transposons (33%), five short interspersed nuclear element (SINEs, 17%), 18 long terminal repeat (LTRs, 28%) and 53 long interspersed nuclear element (LINE, 68%) families (**Fig. 1b**). Interestingly, when analyzed by TE class, only LTR elements showed significant up-regulation, a finding which again replicated in both the Matched and Unmatched PD groups (**Additional File 1: Figure S1b**). These results suggest that loss of TE repression, and particularly LTR elements, may play a role in PD pathogenesis.

Focussing on dysregulated LTRs in Matched PD, we found that the majority of differentially regulated elements were from the ERVL and ERVL-MaLR families, with MLT2B3, LTR70, MLT1H1, and THE1D showing both high average expression and up-regulation. LTR12B elements (ERV1 sub-family) were the most expressed and up-regulated (**Fig. 1c**), a finding that again replicated in Unmatched PD samples (**Additional File 1: Figure S1c**). LTR12B therefore was the most abundant and up-regulated ERV1 in PD samples. Of all investigated LINE elements, only those belonging to the L1 family showed differential expression. L1s showed the highest average expression in both Matched and Unmatched PD samples to controls (**Fig. 1d**, **Additional File 1: Figure S1d**).

We next tested for evidence of co-regulation between TEs and genes. We performed gene set enrichment analysis (GSEA), and in a second step, correlated the GSEA results with DETE dysregulation. Of > 7,000 pathways investigated, we found 213 up- and 608 down-regulated in Matched PD relative to controls (FDR q < 0.05; **Additional File 3**). Intriguingly, the enrichments included up-regulated clusters of immune response pathways including ‘positive regulation of immune response’ (GO:0050778), ‘response to external biotic stimulus’ (GO:0043207), and ‘diseases associated with TLR signaling cascade’ (R-HSA-5602358.1) (**Fig. 1e**), pathways whose activation has been associated with exogenous and endogenous viral activation. Consistent with our previous results, parallel analysis of Unmatched PD samples showed a highly overlapping set of pathway dysregulation, once again with more modest enrichment scores (**Additional File 3**). Similar to the Matched-PD analysis, the enriched pathways included clusters associated with defense response and inflammation (**Additional File 1: Figure S1e**). While the ‘defense response to virus’ (GO:0051607) was up-regulated, ‘viral life cycle’ (GO:0019058) and ‘viral process’ (GO:0016032) pathways were both down-regulated in Matched PD analysis. Collectively, these results suggested that increased TE expression may induce viral mimicry responses and inflammation in the PD brain.

Next, for each TE and cluster of pathways in Fig. 1e we fit a linear mixed effects model with gene expression as the response variable. Intriguingly, with the exception of LTR41, TEs had negative associations with pathway expression (FDR q < 0.05; **Fig. 1f**). Further, a highly co-varying matrix of TE-to-pathway regulation was observed for those pathways down-regulated in PD. Specifically, pathways associated with respiration and metabolism, HIV, macroautophagy, neuronal projection development, synaptic transmission, and translation were inversely associated with a large number of TEs (**Fig. 1f**).

Thus, TE activation or misexpression is a novel genomic feature of both mild (Unmatched) and progressed (Matched) PFC pathogenesis in human PD. The identified TE-dysregulation-associated gene expression changes support PD pathogenesis models that include viral mimicry and infection-based triggering in PD etiology.

### TEs are up-regulated in sorted PFC neuronal nuclei in PD

To gain insight into the cellular origin of the TE expression signature, we performed RNA-seq of sorted neurons of PFC brain tissue samples of the same Matched (N = 13) and Unmatched (N = 11) PD samples and healthy controls (N = 14). We obtained 15,871 genes and 150 TEs reliably quantified across all samples. After normalization and variance correction we applied a robust linear regression model to each gene with condition as the dependent variable. We adjusted the models for patient age, sex, post-mortem interval, brain hemisphere and possible sources of unknown unwanted variation estimated using RUV.

Comparing Matched PD samples to controls, we identified 851 DEGs and 6 DETEs (**Additional File 4**). DEGs skewed towards up-regulation (OR = 1.75, p = 4.5 × 10^−14^, Fisher’s exact test), and, consistent with the observations made in bulk PFC tissue, all DETEs were up-regulated (**Fig. 2a**). Notably, whereas the Unmatched PD sample analysis validated a DEG skew towards up-regulation (OR = 1.36, p = 0.003), no statistically significant up-regulation of TE expression was detected (OR = 3.51, p = 0.4, Fisher’s exact test). Importantly, analysis comparing Matched and Unmatched PD differential expression showed a very strong correlation for both DEGs and DETEs (Pearson r = 0.68, p < 2.2 × 10^−22^ and r = 0.57, p = 1.57 × 10^−14^ respectively) indicating high similarity between the transcriptional dysregulation in the two conditions. Consistent with the idea that contralateral pathology tends to be more progressed relative to the ipsilateral hemisphere, DEGs were more strongly dysregulated when comparing Matched to Controls than it was when comparing Unmatched to Controls (mean difference 0.12, p = 3.94 × 10^−110^ and 0.26, p = 6.74 × 10^−5^ among the up-regulated genes and TEs, respectively, and mean difference −0.15, p = 9.32 × 10^−112^ and −0.21, p = 0.04 among the down-regulated genes and TEs, respectively, paired t-test, **Additional File 1: Figure S2a**).

**Fig. 2:**
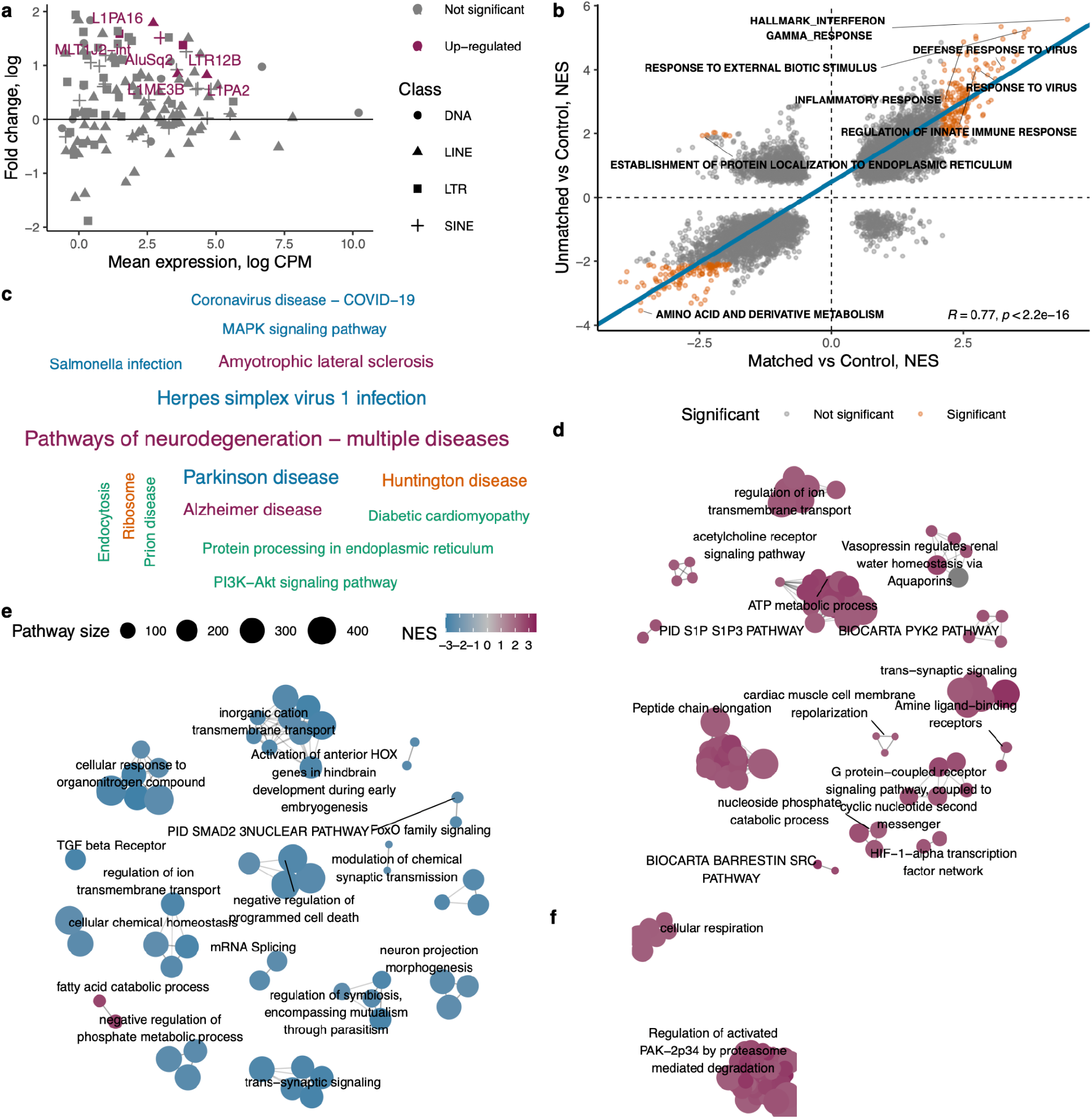
Differential expression of genes and up-regulation of transposable elements (TEs) in sorted prefrontal cortex neurons comparing Matched PD to control brain samples. (**a**) Mean expression and fold change of TEs colored by statistical significance and shaped by element class. **(b)** Correlation of normalized enrichment scores obtained through GSEA of Matched/Control and Unmatched/Control comparisons. Pathways, significantly enriched in both comparisons are shown as orange dots. Selected pathways have been labeled. The blue line was plotted using deming regression coefficients. (**c**) A word cloud of significantly enriched KEGG pathways obtained through GSEA of Matched/Control comparison. (**d**) Network and clustering of the significantly enriched pathways obtained through GSEA of Matched/Control comparison. (**e**) Network and clustering of the significantly enriched pathways obtained through GSEA of genes correlating with MLT1J2. (**f**) Network and clustering of the significantly enriched pathways obtained through GSEA of genes correlating with LTR12B. GSEA, gene set enrichment analysis.

As with the whole PFC tissue analysis we performed GSEA for both the Matched and Unmatched PD analyses. We tested for correlation of normalized pathway enrichment scores in Matched vs Controls and Unmatched vs Controls and found it to be significant (Pearson correlation r = 0.77, p < 2.2 × 10^−16^ and overlap of significantly enriched pathways OR = 5.77, p < 2.2 × 10^−16^; Fisher’s exact test; **Fig. 2b**). Neurodegeneration, Parkinson’s and Alzheimer’s disease as well as Herpes simplex virus and salmonella infection pathways as defined in the KEGG database were significantly enriched (**Fig. 2c**; and **Additional File 1: Figure S2b** for Unmatched). Using enrichment map gene sets from Bader lab and clustering the top 50 most dysregulated pathways we identified multiple PD-related clusters (**Fig. 2d**). In addition to a strong set of enrichments for cell-signaling cascades including acetylcholine and GPCR signaling, several viral pathways including viral mRNA translation (Reactome pathway R-HSA-192823.3) were also observed (**Additional File 5** and **Additional File 1: Figure S2c** for Unmatched). Thus, PFC neurons exhibit progressive TE derepression.

Within the large set of DETEs, MLT1J2-int and LTR12B were both differentially expressed in bulk and sorted neuron data. The expression of LTR12B in sorted neuronal data also correlated with that in bulk tissue (Pearson r = 0.78, p = 2.1 × 10^−8^). These data suggested that LTR12B expression changes in bulk tissue are driven by the neuronal compartment of that tissue. We therefore asked whether the expression of LTR12B or MLT1J2 might be associated with the expression of genes in neuronal tissue. Using linear regression we identified N = 328 genes correlated with expression of LTR12B and N = 184 genes correlated with expression of MLT1J2 (robust linear regression adjusted for condition, brain hemisphere, age, sex, PMI and two RUV vectors representing unwanted variation; **Additional File 6**). Dataset intersection revealed 14 genes that correlated with both of these TEs, an overlap that exceeded random expectation (OR = 4.44, p = 1.04 × 10^−5^) and an analysis that suggested these were acting in concert (Pearson r = 0.82, p = 3.6 × 10^−4^). Intriguingly, the genes included NEURL1, HHIP-AS1, VWC2, ARLNC1, MIR4500HG, PMS2P3, SMARCE1P5, ANOS1, LRRIQ1, and NIPA2, many of which have previously been associated with PD and neurodegeneration^22–26^. Interestingly, MTL1J2 was primarily associated with down-regulated pathway signatures including signaling and negative regulation of cell death (**Fig. 2e, Additional File 7**). LTR12B was positively associated with pathways pertaining to cellular respiration and regulation of PAK-2p34 by proteosome mediated degradation (**Fig. 2f, Additional File 7**). These findings suggest that activated HERV and LINE element expression might trigger or modulate apoptosis in the PD brain.

### TEs are up-regulated and dynamic in peripheral blood of PD patients

The data above identified TE dysregulation as a potential new feature of PD pathology and one that exhibits strong associations with viral mimicry and infection-like gene expression signatures. Substantial evidence in the literature supports the hypothesis that PD is originally triggered in the periphery through infection. We therefore tested whether similar associations were measurable in the patient periphery in PD (ie. outside the CNS). We analyzed whole blood RNA-seq data provided by the Parkinson’s Progression Markers Initiative (PPMI) and evaluated gene and transposable element expression differences between patients with PD (N = 324), Prodromal patients with PD (N = 29) and healthy, age-matched control individuals (N = 167) of Caucasian race, with a mean age of 62.38 ± 10.04 SD. We analyzed data from samples obtained longitudinally, over five visits (0, 6, 12, 24, 36 months), starting from the date of disease diagnosis (0 months). After QC and preprocessing we obtained 17,105 genes and 135 TE reliably quantified across all samples. As above, we applied a multivariate linear regression model after normalization and variance correction, correcting for visit, age, and sex as well as technical parameters (sample position on the plate, sample collection phase, center number) and unknown sources variation.

Comparisons of whole blood transcriptomes of PD and control samples identified 3,497 DEGs and 21 DETEs (FDR q < 0.05, least squares linear regression, **Additional File 8**). Intriguingly, both genes and TEs were once again skewed towards up-regulation (OR = 1.68 and OR = 2.65 respectively; **Fig. 3a**). Among the 21 DETEs, 19 were up-regulated (**Fig. 3b**). Importantly, and in keeping with the PFC analysis in **Fig. 1**, we observed up-regulated TEs in each class, and binomial regression indicated that this association was again driven by LTR transposable elements.

**Fig. 3.**
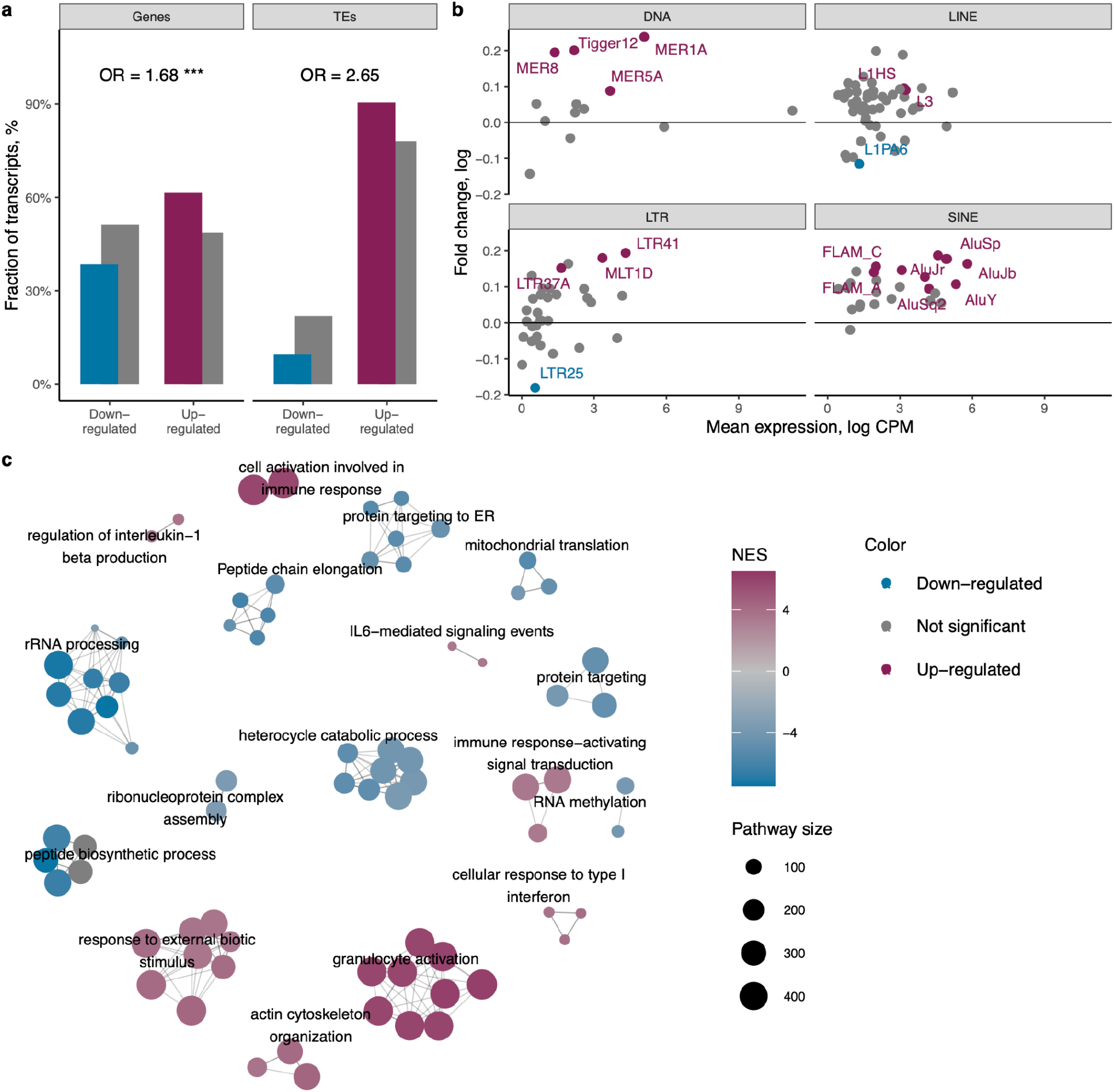
Differential expression of genes and transposable elements (TEs) in blood of Parkinson’s disease patients. **(a)** Fraction of up-regulated and down-regulated significant genes and TEs (colored fill) compared to the fraction of non-significant up-regulated and down-regulated genes as well as TEs (gray fill). **(b)** Mean expression and fold change of TEs stratified by class and colored by statistical significance. **(c)** Network and clustering of the top 50 most up-regulated and 50 most down-regulated pathways obtained through gene set enrichment analysis. Each cluster is named after the most prominent pathway decided by the PageRank algorithm.

GSEA analysis identified 863 up-regulated and 254 down-regulated pathways at FDR q < 0.05 when comparing PD to Control samples (**Additional File 9**). Clustering of the top 50 most significantly up-regulated and 50 most down-regulated pathways revealed DNA replication, RNA modification, and mitochondrial translation clusters down-regulated in PD, while oxidative stress, mitochondrial function and numerous immune response pathways including IL6, IL1b and type-1 interferon (viral) response pathways were up-regulated (**Fig 3c**). Together with the findings above, these data demonstrate hallmark features of PD transcriptional dysregulation found in the brain are already present in peripheral blood as early diagnosis. They support PD pathogenesis models where PD is considered a systemic inflammatory disease ^27^.

Comparisons of Prodromal PD with Controls at baseline visit identified substantially fewer DEG’s (1,700) and 14 DETEs (FDR q < 0.05; least squares linear regression). Once again DEG’s trended towards transcriptional up-regulation (OR = 1.09, p = 0.11; **Additional File 1: Figure S3a**; **Additional File 8**). From a statistical standpoint, it is noteworthy that in addition to milder disease severity in the Prodromal individuals, the sample number was much smaller than the PD group. Interestingly, TEs in Prodromal PD were down-regulated relative to Controls despite a strong positive correlation in both differential gene expression (Pearson r = 0.29, p < 2.2 × 10^−16^; **Additional File 1: Figure S3b**) and DETEs (Pearson r = 0.28, p = 0.001; **Additional File 1: Figure S3c**) when comparing the PD and Prodromal PD responses to Control. Normalized Enrichment Scores between the two conditions showed substantial overlap of significant pathways (r = 0.33, p < 2.2 × 10^−16^ and OR = 4.59, p < 2.2 × 10^−16^; Pearson correlation and Fisher’s exact test) though it is noteworthy that the co-regulation amongst the top 100 differentially regulated pathways were substantially different (**Figure S3d**). Thus, while central transcriptional features of PD can be observed in the periphery as early as diagnosis, from a blood transcriptome point of view the prodromal PD is distinct from post-diagnosis.

Since TE up-regulation appeared to be triggered at or around PD diagnosis, we performed a dedicated longitudinal analysis. The analysis revealed 6,058 DEGs (FDR < 0.05) and 66 DETEs associated in some way with PD over time (**Additional File 8**). Unexpectedly, we observed a significant down-regulation of both genes and TEs with time (OR = 0.88, p = 5.56 × 10^−5^ and OR = 0.07, p = 1.59 × 10^−5^ for genes and TEs, respectively; Fisher’s exact test; **Additional File 1: Figure S4a**), a finding reciprocal to the baseline comparison of PD samples against Controls. Thus, blood transcriptome activation is a hallmark feature of PD that attenuates over time.

Given that the PPMI sample set zero time point is the time of diagnosis, we hypothesized that the attenuation of these inflammatory signatures might be linked to the initiation of PD treatment. A comparison between PD-associated gene expression and the genes that change progressively over time in PD revealed a strong correlation indicating that PD is also a progressive disease from the perspective of the blood transcriptome (r = 0.37, OR = 1.47, p < 2.2 × 10^−16^; **Additional File 1: Figure S4b**). Noteworthy, DETEs exhibited the opposite association, if any (OR = 0.2; **Additional File 1: Figure S4c**). Indeed, analysis of TE dynamics over time showed a significant association between reduced transposable element expression and its significance in PD over time (p = 3.91 × 10^−6^, binomial regression,), a statistical effect that was significantly driven by LTR TEs (p = 5.66 × 10^−5^, binomial regression; **Additional File 1: Figure S4d**).

Altogether, the collective PD transcriptional analyses identify PFC TE up-regulation as a novel feature of PD; they indicate that TE up-regulation is triggered in the periphery at or around disease onset; that TE up-regulation ceases and in some cases reverses upon diagnosis and treatment onset; and, the data identify LTR TEs as potential novel mediators or readouts of PD medication efficacy and compliance.

### A Trim28/ERV axis buffers against variability in PFF-triggered pathology

Finally, we aimed to test whether TE up-regulation might play a causal role in PD pathogenesis. We generated mice harboring a heterozygous loss-of-function mutation in the TE silencing factor Trim28 (Trim28^+/D9^) and challenged those animals with a PD-mimicking alpha-synuclein preformed fibril (PFF) injections into the dorsal striatum. The experimental groups consisted of the two genotypes (WT and Trim28^+/D9^) for which groups of sham-treatment (PBS) and PFF injection were both run. Animals were euthanized after one month to avoid indirect or secondary effects. In addition to histopathology, RNA-seq was performed on a total of N = 38 substantia nigra brain samples.

We first performed a baseline characterization of the model. After QC and preprocessing we used multivariate linear regression to adjust for age, sex, lean and fat mass as well as possible sources of unknown variation. RNA-seq confirmed down-regulation of Trim28 gene expression in Trim28^+/D9^ animals (**Additional File 1: Figure S5a**). Consistent with the role of Trim28 as a repressor of TE expression, Trim28^+/D9^ animals showed modest up-regulation of TEs at baseline (**Fig. 4a; Additional File 11**). Similar to our observations in PD patient neuronal and blood datasets, TE up-regulation was accompanied by a skew towards genic down-regulation (OR = 0.67, p = 9.7×10^−8^ and OR = Inf, p = 0.08 for genes and TEs, respectively; Fisher’s exact test; **Fig. 4a**). Five of the 6 significantly dysregulated TEs were ERVs of LTR class which is consistent with Trim28 literature (**Fig. 4b**). Intriguingly given the lack of obvious behavioral phenotypes in most Trim28^+/D9^ mice, GSEA identified 172 up-regulated and 1,181 down-regulated pathways (FDR q < 0.05; **Additional File 12).** Cytokine signaling in the immune system, cell recruitment (pro-inflammatory response) pathway clusters as well as viral infection KEGG pathways were enriched (FDR q < 0.05; **Additional File 1: Figure S5b,c**). Immunohistological examination of sham-operated animals showed no evidence of pathology in either Trim28^+/D9^ or WT animals (PBS groups; **Fig. 4c**). Thus, Trim28^+/D9^ mice exhibit PD-like precocious TE activation and related gene expression effects.

**Figure 4.**
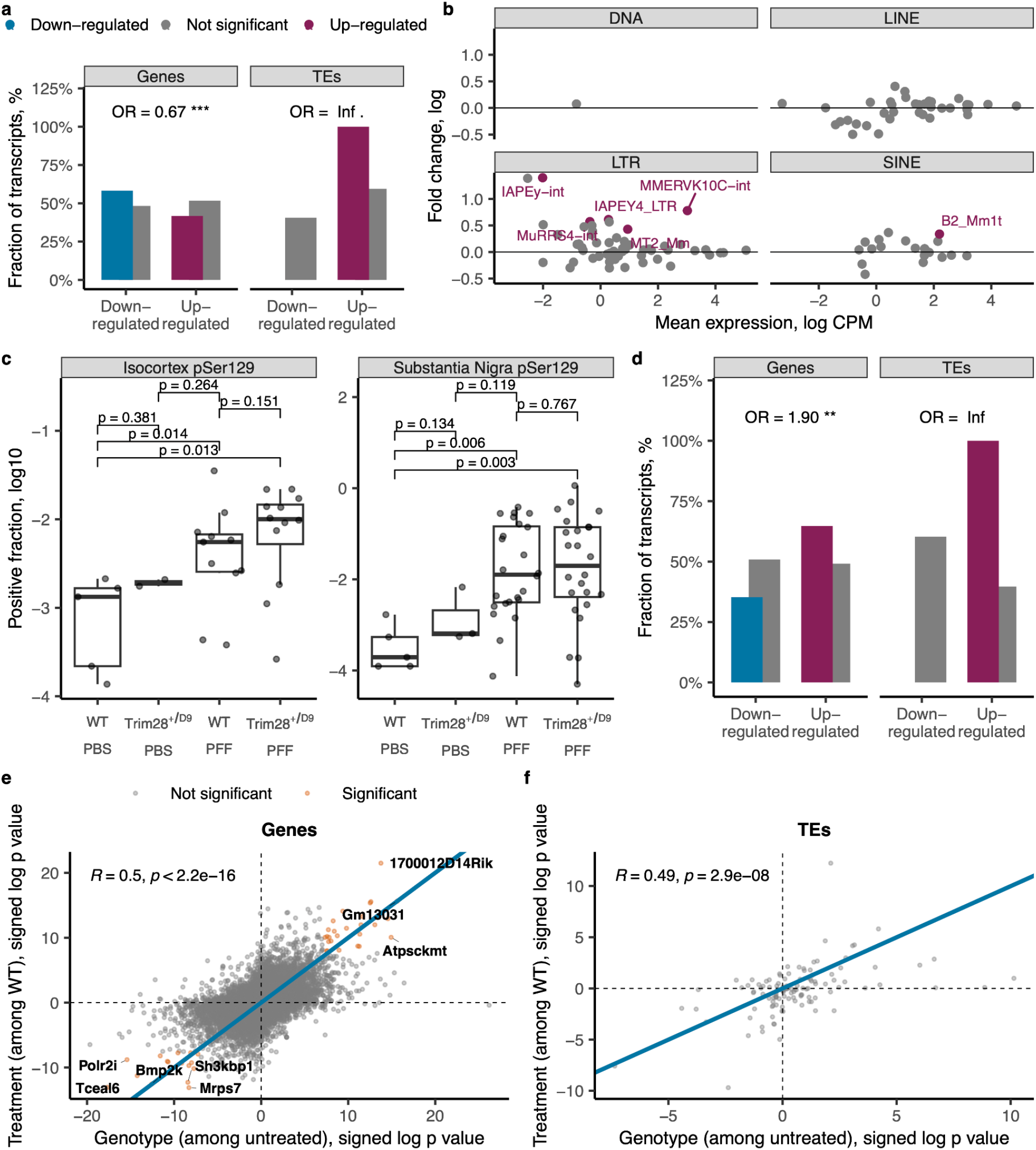
Differential expression of genes and transposable elements (TE) in PFF injected and Trim28 heterozygous knockout mouse substantia nigra samples. (**a**) Fraction of up-regulated and down-regulated significant genes and TEs (colored fill) compared to the fraction of non-significant up-regulated and down-regulated genes as well as TEs (gray fill) in Trim28 heterozygous knockout mouse brain data. (**b**) Mean expression and fold change of TEs stratified by class and colored by statistical significance in Trim28 heterozygous knockout mouse brain data. **(c)** Misfolded alpha synuclein pathology scores (pSER129) in substantia nigra and isocortex tissues. **(d)** Fraction of up-regulated and down-regulated significant genes and TEs (colored fill) compared to the fraction of non-significant up-regulated and down-regulated genes as well as TEs (gray fill) in PFF injected mouse brain data. **(e)** Correlation of gene expression changes in Trim28^+/D9^ to wild-type comparison and PFF injected to PBS injected animal comparison. Significant differentially expressed genes in both comparisons are indicated in orange. Labeled genes at the perimeter indicate the strongest effect in both datasets. Blue line was plotted using deming regression coefficients. **(f)** Correlation of TE expression changes in Trim28^+/D9^ to wild-type comparison and PFF injected to PBS injected animal comparison. Blue line was plotted using deming regression coefficients.

Next, we examined the effect of PFF-injection itself. We observed significantly increased misfolded alpha synuclein pathology scores (pSER129) in isolated substantia nigra and isocortex tissues for both WT and Trim28 heterozygous mice injected with PFF as compared to PBS injected WT controls (**Fig. 4c; Additional File 10**). The latter was expected given the low dose applied, the early time point used and the small fraction of each sample that displayed substantial pathology. While the trend was non-significant, Trim28^+/D9^ animals injected with PFFs showed the greatest level of isocortex pathology (**Fig. 4c**), and potentially interesting given previous work on Trim28, the greatest variability in pathological scores. From the genomic perspective, PFF injection alone showed modest effects on gene expression relative to the effect of the Trim28^+/D9^ mutation with 105 differentially expressed genes and a skew towards up-regulation (OR = 1.90, p = 0.002; Fisher’s exact test; **Fig. 4d; Additional File 11**); and up-regulation of 1 TE (**Additional File 11)**, a LINE L1 subclass TE (**Additional File 1: Figure S5d**). Importantly, gene and TE expression changes induced by PFF injection and Trim28 knockout were highly correlated (Pearson correlation r = 0.5, p < 2.2 × 10^−16^ and r = 0.49, p = 2.9 × 10^−8^, respectively; **Fig. 4e,f**). Early PFF-induced pathology triggers transcriptional changes similar to those triggered by reduced Trim28-dependent silencing, including TE up-regulation. Collectively, these data are consistent with a two-compartment model where genotype generates a widespread genomic predisposition and a second event (in this case PFFs) triggers pathology.

Finally, to understand whether Trim28^+/D9^ triggered ERV up-regulation activates similar pathways to those observed in human PD samples, we tested for overlap in dysregulated gene sets across experiments. From the 4,997 pathways analyzed in both mice and bulk human brain tissue we identified 160 pathways that were significantly enriched in both datasets, a rate significantly higher than expected by chance (OR = 2.16, p = 1.32 × 10^−12^; Fisher’s exact test). We found 71 pathways that were significantly down-regulated in both Trim28^+/D9^ mouse and contralaterally Matched PD samples (OR = 3.73, p = 3.2 × 10^−16^; Fisher’s exact test). Consistent with human analyses, numerous viral response pathways including ‘viral process’ (GO:0016032), ‘viral life cycle’ (GO:0019058), and ‘regulation of apoptosis’ (R-HSA−169911.2) were among the down-regulated pathways (**Fig. 5**). Thus, the coding and non-coding genome regulation observed in human PD shows a significant similarity to the effects of partial Trim28 loss-of-function.

**Figure 5.**
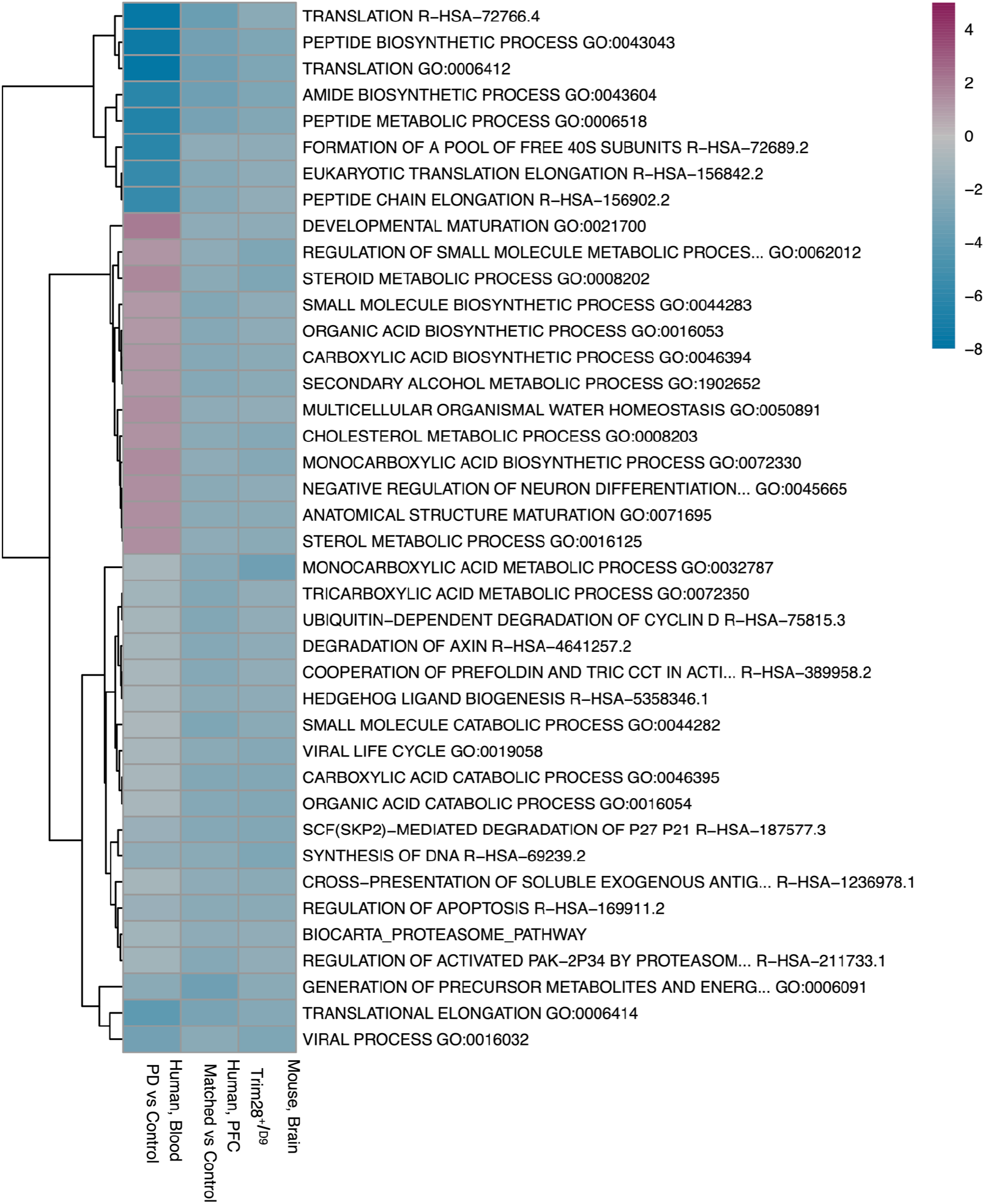
Overlap of enriched PD related pathways in human blood, human prefrontal cortex bulk tissue samples and Trim28 heterozygous knockout mouse brain samples. The fill color indicates signed log p value of pathway enrichment. PD vs control contrast was used for blood data. Matched versus control contrast was used for prefrontal cortex tissue data. Knockout vs wild type contrast was used for mouse brain tissue data.

## DISCUSSION

Our study identifies transcriptional activation of TEs as a hallmark of PD and provides a systematic characterization of the landscape of transcriptional dysregulation in PD prefrontal cortex and in blood. The data demonstrate that intrastriatal injections of aSyn PFFs can trigger genome-wide transcriptional dysregulation in the striatum that mimics partial loss of Trim28, suggesting a potential role for destabilization of ERV repression in PD. These data add to previous work in the field showing that Trim28 is necessary for normal aSyn and Tau expression levels ^28,29^ and that loss of Trim28 during neurodevelopment triggers neuroinflammation ^30^. Collectively, our data together suggest a model where Trim28-dependent suppression of TE expression protects from neuroinflammation. Importantly, our comparative analysis of human PPMI resources with the Trim28^+/D9^ (to mimic physiological downregulation) shows that PD shares similarities with a Trim28 haploinsufficient state.

Our findings have four main implications important for understanding PD. First, we find that expression of LINEs and ERVs is up-regulated in prefrontal cortex tissue and in the blood of advanced stage PD patients. The data show consistent up-regulation of TEs and that this signature is more pronounced on the side of the brain associated with disease symptom onset. TE upregulation is less evident among prodromal patients. The data indicate that TE dysregulation is a dynamic and common feature of PD. Aberrant expression of LINE and endogenous retrovirus (ERV) elements were the most prominent, though variation in TE family dysregulation was observed across tissue types. PD therefore joins a growing group of neuronal diseases whose severity correlates with increased ERV expression ^31–42^. HERV expression can be re-activated in a chromatin and sequence-specific manner during infection and aging ^43–45^ and identifying the relevant mechanistic intermediaries represents an important next step for the field. Studies in multiple sclerosis, Alzheimer’s disease, schizophrenia, amyotrophic lateral sclerosis, and PD have found ERV expression in the cerebrospinal fluid in post-mortem brain biopsies too. Our work indicates that ERV expression dynamics are coupled to events that coincide with diagnosis and/or treatment. The potential of ERVs as biomarkers should therefore be explored. While the downstream effects of ERVs in neurological disorders remains enigmatic, there is evidence that activation of the innate immune response is involved^9,41^.

The data also identify an association between precocious ERV expression and cell death in neuronal tissue. We demonstrate enrichment of pathways related to immune response, response to viral infection in both bulk and sorted neuron tissues as well as in blood, pathways expected to exacerbate inflammatory PD phenotypes. We show that genes correlating with ERV expression are enriched in apoptosis related pathways and associated up-regulation of cellular respiration. We hypothesize that neuroinflammation and cell death is triggered, at least in part, by an innate immune response to aberrant HERV activation. Interestingly, the occurrence of PD is known to be inversely associated with cancer in epidemiological studies ^46^, and retrotransposon activation has been shown to induce cancer resistance through cell death regulation, at least in select contexts ^47^. It will be important to test in PD models whether ERV expression represents a double-edged sword.

Finally, we find evidence that synucleinopathy (PFF injection) and ERV up-regulation elicit overlapping responses in murine experimental PD. Comparing transcriptional signatures of Trim28 haploinsufficiency and PFF injection, we identified substantial similarity in the transcriptional responses and showed these to be enriched for inflammatory expression patterns. PFF injection in wild-type mice alone, however, did not elicit an obvious effect on ERV expression. Taking into consideration the very small fraction of any given sample (tissue section) that contains pathology in those assays, it remains unclear from our data whether PFFs are sufficient to generate an ERV response.

Since Trim28 haploinsufficiency leads to ERV derepression and since Trim28 knockout mice do not exhibit PD phenotypes on their own, the data would suggest that it is the ERV activation, and not other PD related processes, that activates viral response and apoptosis pathways in PD. Given the association between ERV up-regulation and PD severity, it will be interesting to see if suppression of ERV expression specifically might slow down neurodegeneration in PD and thus provide therapeutic benefit. Indeed, anti-inflammatory and antiviral drugs have already been shown through epidemiological studies to be associated with reduced PD incidence ^48–50^.

### Contributions

AB performed the mouse experiments.

JG, MM, LM performed computational and statistical analysis

TG wrote the initial version of the manuscript with input from JG and LM.

All authors contributed to manuscript preparation.

Elizabeth Van Putten performed neuronal nuclei and RNA isolation of human brain samples.

### Data and code availability

Raw data and code used to process the data and obtain the results are available from the authors upon request.

## Acknowledgements

We would like to acknowledge the discussions and help with manuscript editing by Migle Gabrielaite.

## METHODS

### Prefrontal Cortex Tissues Samples

Prefrontal cortex tissue were obtained from the Parkinson’s UK Brain Bank, NIH Neuro-BioBank, and Michigan Brain Bank, with approval from the ethics committee of the Van Andel Research Institute (IRB #15025). For each individual, we have information on demographics (age and sex assigned at birth), tissue quality (postmortem interval, tissue quality/RIN score), and clinical variables (PD diagnosis), provided in **Additional File 13**. Control individuals had pathologically normal brains (and verified to have no brain Lewy body pathology), while PD cases were pathologically confirmed to have brain Lewy body pathology. Neurons of the prefrontal cortex were examined in this study because 1) this brain region plays an important role in PD ^51^, 2) pathology does spread to this brain region ^52^, and 3) prefrontal cortex neurons still exist in the postmortem PD brain ^53^. Neuronal nuclei isolation from prefrontal cortex tissue was done by a flow cytometry-based approach, as described ^54^.

### Prefrontal Cortex RNA Sequencing Libraries

We used RNA-seq to profile the gene and TE transcriptome in both bulk and neuronal prefrontal cortex samples from PD and healthy controls. Total RNA was isolated using the standard TRIzol protocol and re-suspended in 85 μL of ultrapure distilled water (Invitrogen). DNase treatment was performed with the RNase-Free DNase kit (Qiagen) using 5 μL DNase I and 10 μL RDD buffer, followed by a column cleanup using the RNeasy Mini kit (Qiagen) with two additional washes (75% ethanol) before elution. RNA quantity was assessed by Nanodrop 8000 (Thermo Scientific) and quality was assessed with an Agilent RNA 6000 Nano Kit on a 2100 Bioanalyzer (Agilent Technologies, Inc.). RNA-seq libraries were prepared by the Van Andel Genomics Core from total RNA using the KAPA RNA HyperPrep Kit with RiboseErase (v1.16) (Kapa Biosystems). Base calling was done by Illumina NextSeq Control Software (NCS; v2.0), and the output of NCS was demultiplexed and converted to FastQ format with Illumina Bcl2fastq (v1.9.0).

### Parkinson’s progression markers initiative (PPMI) Samples

This study was done in collaboration with the Parkinson’s progression marker initiative (PPMI). PPMI provided RNA sequencing data from whole blood of PD and healthy control patients unaffected by neurologic diseases. The PPMI cohort contains multiple samples of the same patients across three years. At enrollment, PD patients were required to be untreated with PD medications (levodopa, dopamine agonists, MAO-B inhibitors, or amantadine), within 2 years of diagnosis, Hoehn and Yahr score of less than three, and to have either at least two of resting tremor, bradykinesia, or rigidity (must have either resting tremor or bradykinesia). PD patients also had to show a dopamine deficit via dopamine transporter (DAT) imaging or vesicular monoamine transporter (VMAT-2) imaging. At enrolment the prodromal patients were required to be over 60 years old, have no clinical diagnosis of PD, parkinsonism or dementia, have REM sleep behavior disorder (RBD) and Hyposmia based on UPSIT testing. Healthy controls at enrolment have no current or active clinically significant neurological disorder, no first-degree relative with PD, and normal dopamine levels via DAT imaging.

### Generation of Trim28 Mice

Trim28^+/D9^ were initially generated during N-ethyl-N-nitrosourea mutagenesis screenings as described in Blewitt et al., 2005 ^55^, described therein as *modifiers of murine metastable epialleles* D9 (MommeD9). The phenotypic variation of this line was further characterized in ^19^, demonstrating that the mutants with aberrant ERV expression in adipose tissue have increased adiposity. Animals were under a 12 hour day/night cycle, and housed based on international guidelines. Food and water administered *ad libitum*.

### Stereotactic injections

Purification of recombinant full-length mouse α-syn fibrils (PFFs) was performed as previously described ^56,57^. On the day of stereotactic injection, PFFs were thawed, sonicated at room temperature by probe sonication (Misonix XL2020, pulse sonication at 50% amplitude with 1 sec on and 1 sec off for 2 minutes of total sonication time) and kept on ice for the duration of the surgical procedures. Animals of 20-22 weeks of age were anesthetized with isoflurane/oxygen mixture and stereotactically injected with either sterile PBS or 1 ug PFFs (1 ug/uL) into the dorsal striatum (coordinates: AP +0.2 mm, L −2.0 mm, DV −2.6 mm relative to bregma and dural surface) with a 10 uL Hamilton microsyringe at a constant rate of 0.2 uL per minute as previously described ^58^. The microsyringe was left in place for 3 minutes after injection, and then slowly removed. This dosage was decided upon after a serial dilution was performed in C57B6J animals and determined to be the lowest amount to induce pathology.

### Tissue preparation

Tissue was collected when animals reached 36 weeks of age. Mice were anesthetized with tribromoethanol and perfused transcardially with 0.9% saline, followed by 4% PFA in phosphate buffer. Brains were harvested, post-fixed for 24 h in 4% PFA at 4°C, and cryoprotected in 30% sucrose in a phosphate buffer for at least 3 days at 4°C or until taken for sectioning. The entire brain of each mouse was cut into 40 µm free-floating coronal sections on a freezing microtome and stored in a cryoprotectant solution consisting of 30% sucrose and 30% ethylene glycol in PBS at −20°C until immunostaining.

### Immunohistochemistry

Coronal free-floating sections were stained with one of the following primary antibodies (xyx). For detection of the antibody with DAB, we utilized a peroxidase-based method (Vectastain ABC kit and DAB kit; Vector Laboratories). Stained sections were mounted onto gelatin-coated glass slides, dehydrated, and coverslipped with Cytoseal 60 mounting medium (Thermo Fisher Scientific). Tyrosine-hydroxylase-stained and mounted sections were counterstained with Cresyl Violet prior to dehydration and coverslipping.

### RNA Sequencing Data Analysis

The same RNA sequencing data analysis pipeline was used for all human and mouse datasets as follows. Adapter trimming, removal of PCR duplicates and quality filtering was performed using fastp (v0.23.2) ^59^. Read alignment and gene counts were performed using STAR (v2.7.9a) ^60^. Reference genome and gencode versions were GRCh38 and 38 for human and GRCm38 and M25 for mice samples, respectively. TE counts were obtained using TEtranscripts (v2.2.1) ^61^. Gene counts matrix was used to find DE genes. Combined gene and TE counts estimated by TEtranscripts were used to identify DETEs. The count matrices were imported into R (4.1) and edgeR (v3.38.1) ^62^ was used to remove low expressed genes prior to trimmed fastmean of M-values normalization. Voom variance stabilization and robust or ordinary least squares regression implemented in limma (v3.50.1) ^63^ was used to identify DE genes and TEs. The regression models were adjusted for known confounders as well as sources of unknown variation, which were determined using stable control genes (p value ≥ 0.5) using the RUVseq Bioconductor package (v1.28.0) ^64^. Multiple testing correction was performed using Benjamini-Hochberg correction.

For the bulk prefrontal cortex samples Cell-type deconvolution was performed using CIBERSORT ^65^ (100 permutations) with 834 brain cell-type-specific gene signatures ^66^. The robust linear regression models were adjusted for diagnosis, age, sex assigned at birth, RIN, hemisphere, and neuronal proportion. For the neuronal prefrontal cortex samples the robust linear regression models were adjusted for diagnosis, age, sex assigned at birth, RIN, brain hemisphere, post-mortem interval, and sources of unknown variation. For the sorted neurons of prefrontal cortex samples robust linear regression models were adjusted for diagnosis, age, sex assigned at birth, brain hemisphere, post mortem interval and sources of unknown variation. Analysis of the whole blood PPMI data was performed with a linear regression model adjusted for the combination of diagnosis and month, sex assigned at birth, age, position, phase, sample processing centre number and sources of unknown variation, with a blocking factor for patient identification.

For the Trim28 heterozygous mice samples the robust linear regression models were adjusted for condition, sex, lean mass at 16 weeks and adjusted fat mass. Condition was a combination of genotype (either WT or heterozygous knockout) and injection (either PFF or PBS). The adjusted fat mass was computed as residuals from a linear model where fat mass was the response variable and genotype and sex were the independent variables.

### Gene set enrichment analysis

Pathway enrichment was performed using GSEA (v4.1.0) software from https://www.gsea-msigdb.org/gsea/index.jsp. Genes were ranked according to the sign of their DE fold change and negative log transformed p value. Mouse and human gene set files (GMT) were downloaded from Bader lab website on August 03 of 2022 ^67^. 1000 permutations were performed to estimate pathway enrichment p values. Fifty of the most significant up-regulated pathways and fifty of the most significant down-regulated pathways were clustered and their interaction networks were visualized using the aPEAR package ^68^. Each cluster was named after the most influential pathway within that cluster determined using the PageRank algorithm. In addition to the above, R package clusterProfiler was used to obtain the word cloud of the top 100 most significant pathways.

## Notes

### Competing Interest Statement

The authors have declared no competing interest.

