## Supplementary figures and images for "Human Endogenous Retrovirus Expression is Dynamically Regulated in Parkinson’s Disease"

### Figure S1

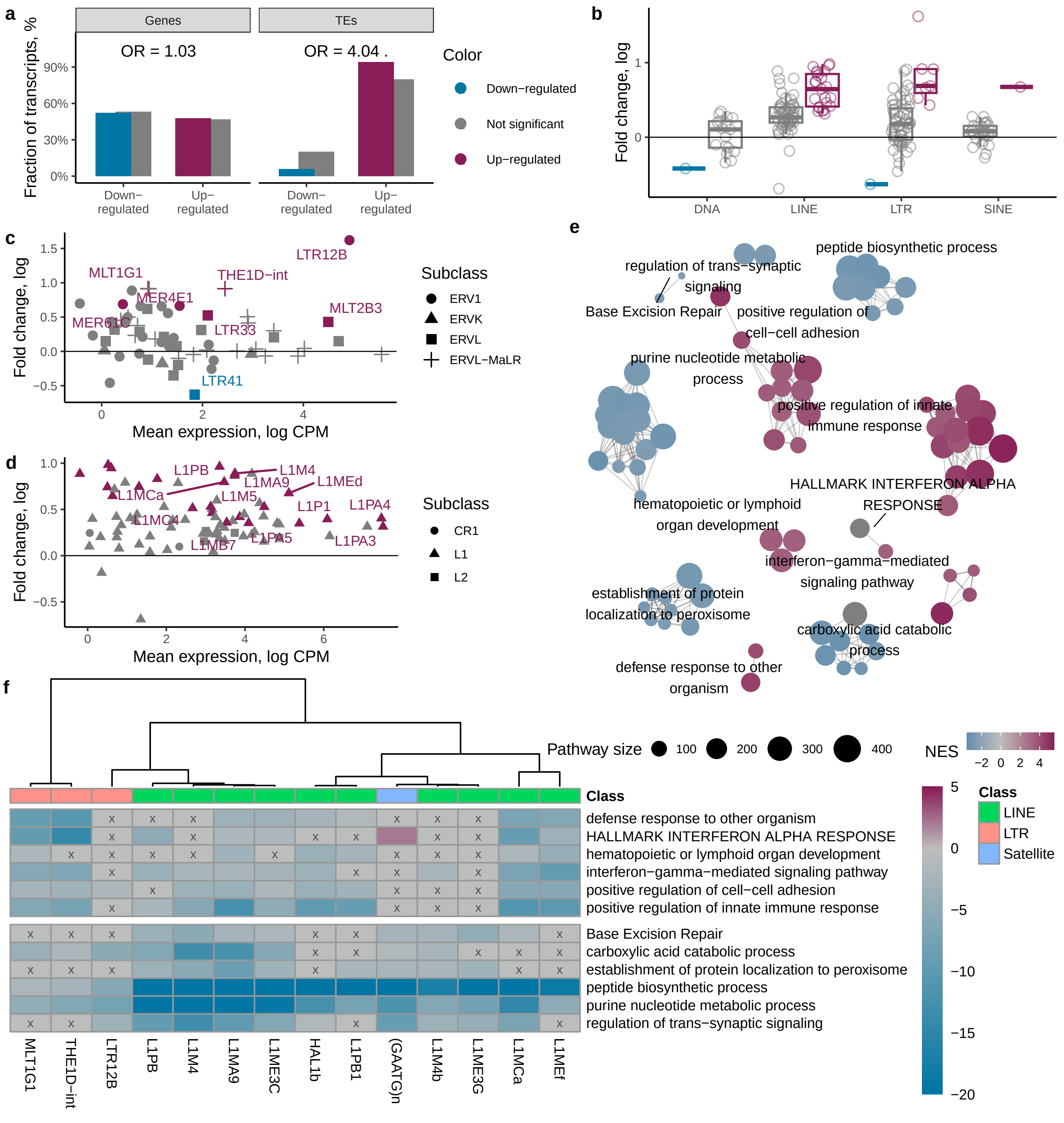

### Figure S2

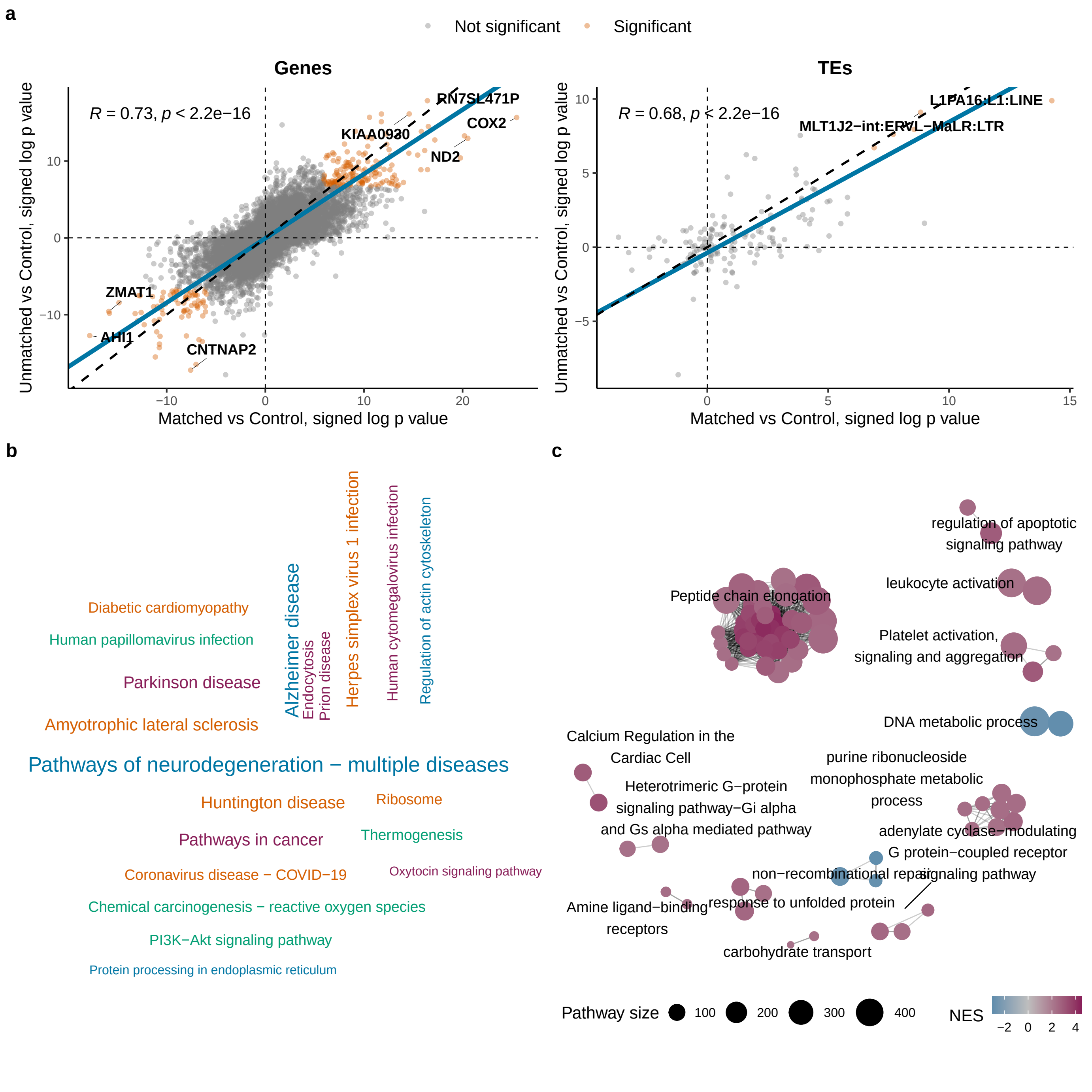

### Figure S3

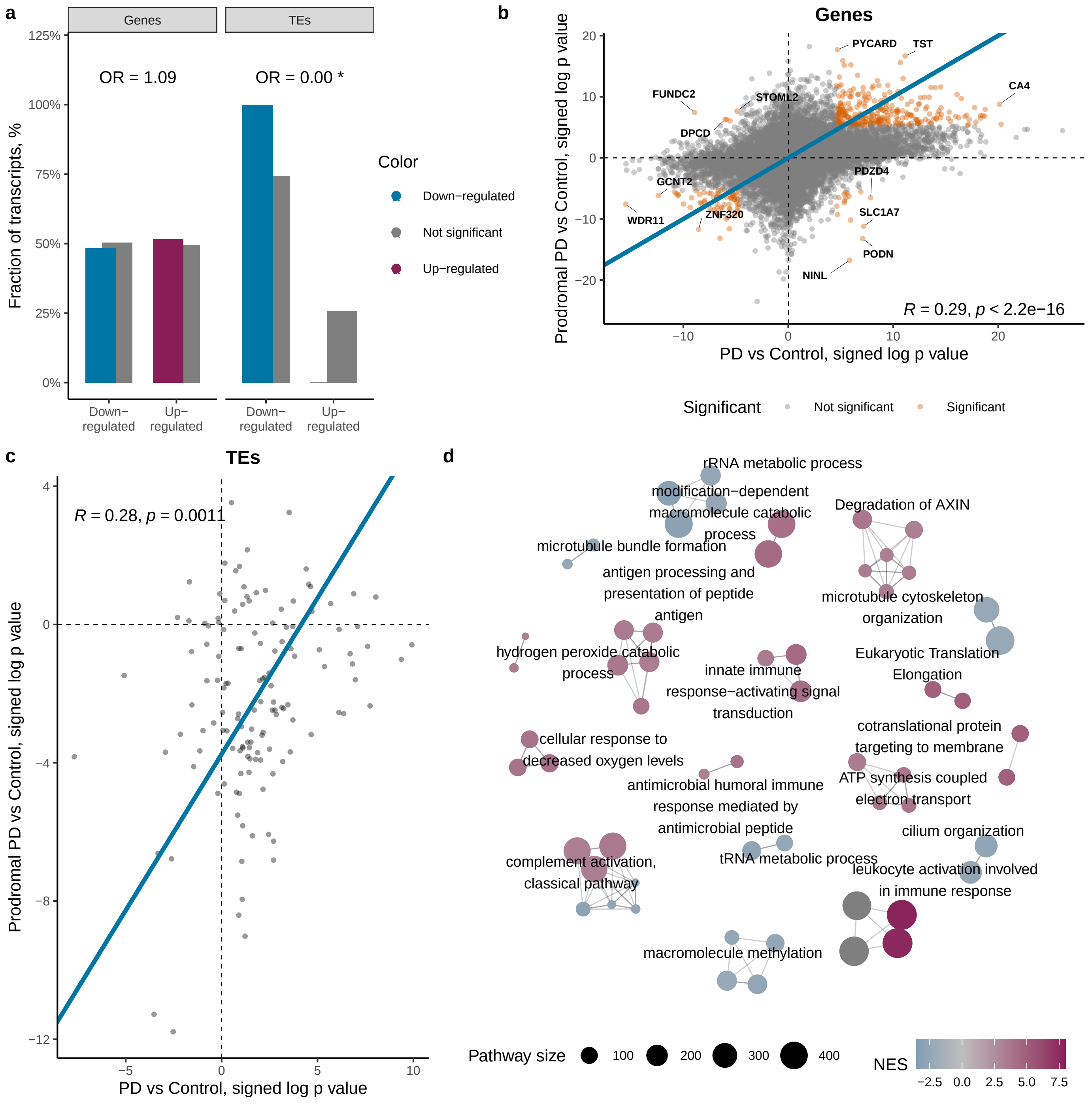

### Figure S4

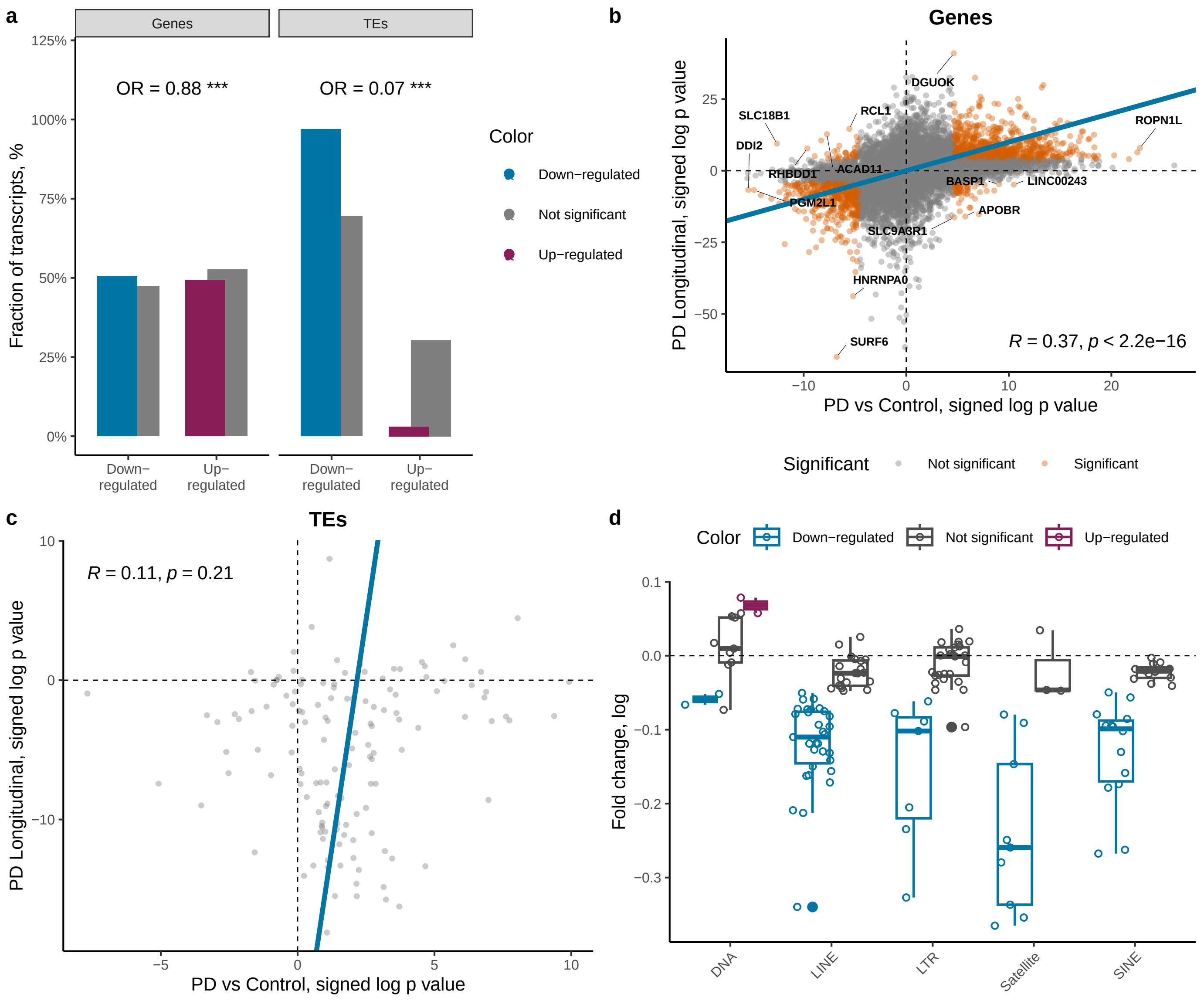

### Figure S5

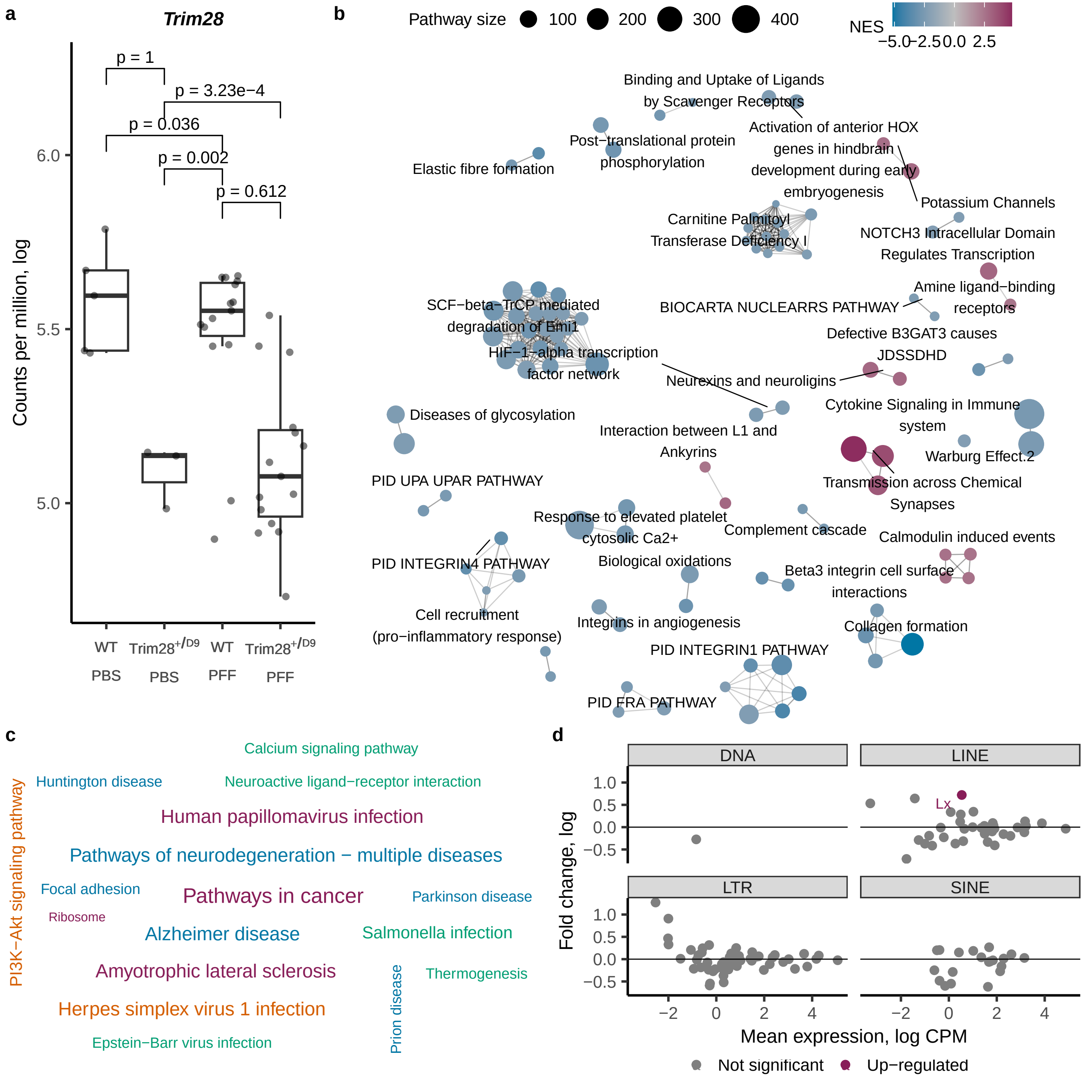
