## Additional File 1 for "Human Endogenous Retrovirus Expression is Dynamically Regulated in Parkinson’s Disease"

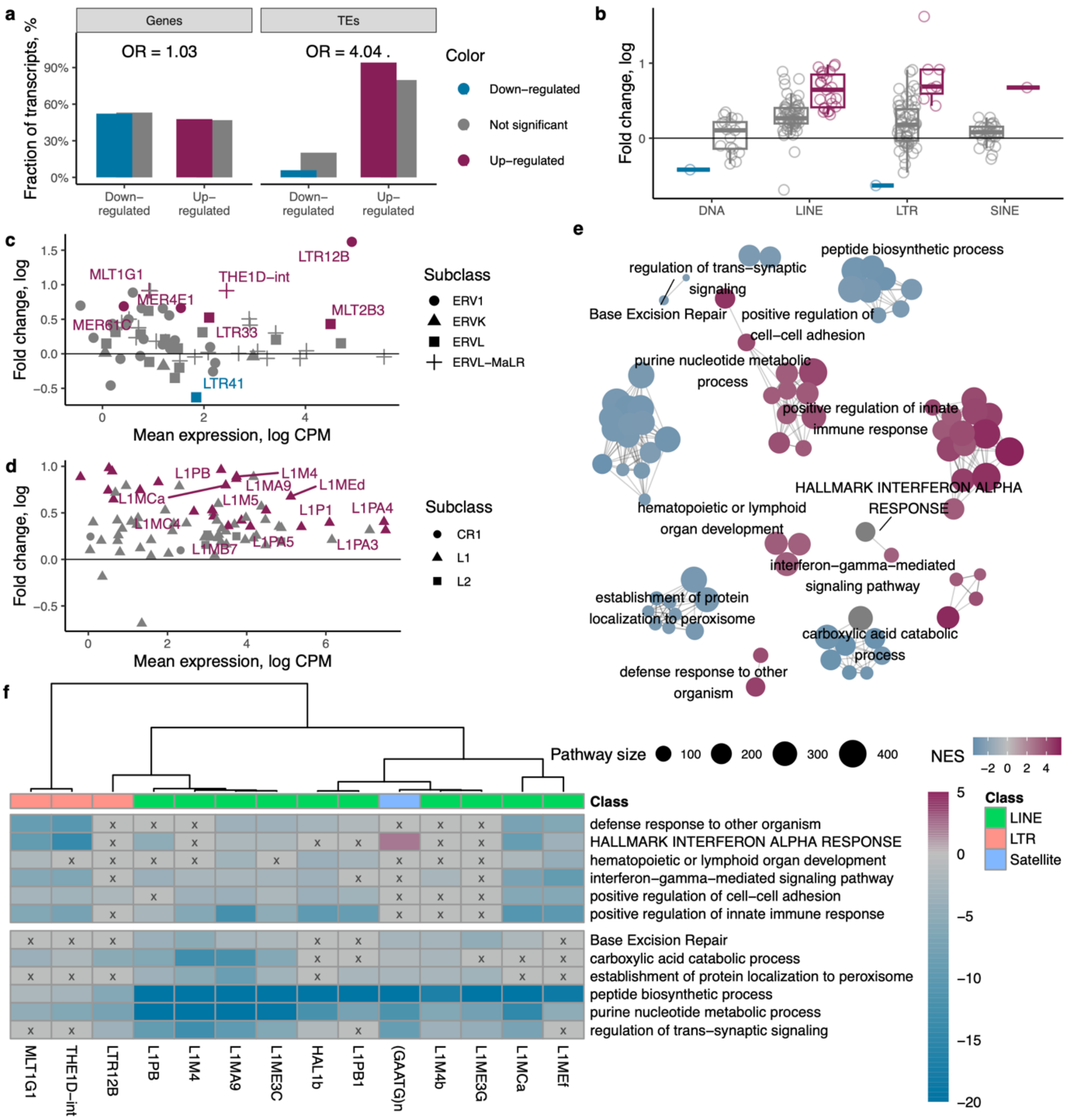

**Figure S1. Differential expression of genes and up-regulation of transposable elements (TEs) in the bulk prefrontal cortex neuron tissue comparing Unmatched PD to control brain samples. (a)** Fraction of up-regulated and down-regulated significant genes and TEs (colored fill) compared to the fraction of non-significant up-regulated and down-regulated genes as well as TEs (gray fill). **(b)** Fold change of TEs stratified by class and colored by statistical significance. **(c)** Mean expression and fold change of long terminal repeat elements (LTRs) colored by statistical significance and shaped by element subclass. **(d)** Mean expression and fold change of LINE elements colored by statistical significance and shaped by element subclass. **(e)** Network and clustering of the top 50 most up-regulated and 50 most down-regulated pathways obtained through gene set enrichment analysis. Each cluster is named after the most prominent pathway decided by the PageRank algorithm. **(f)** Mixed effects modeling of the association between the expression of TEs and genes belonging to the leading pathways indicated in clustering analysis. Fill color indicates signed log p value. P values were adjusted for multiple testing using FDR. Heatmap cells with a value exceeding FDR q-value higher than 0.05 were marked with an x and gray color.

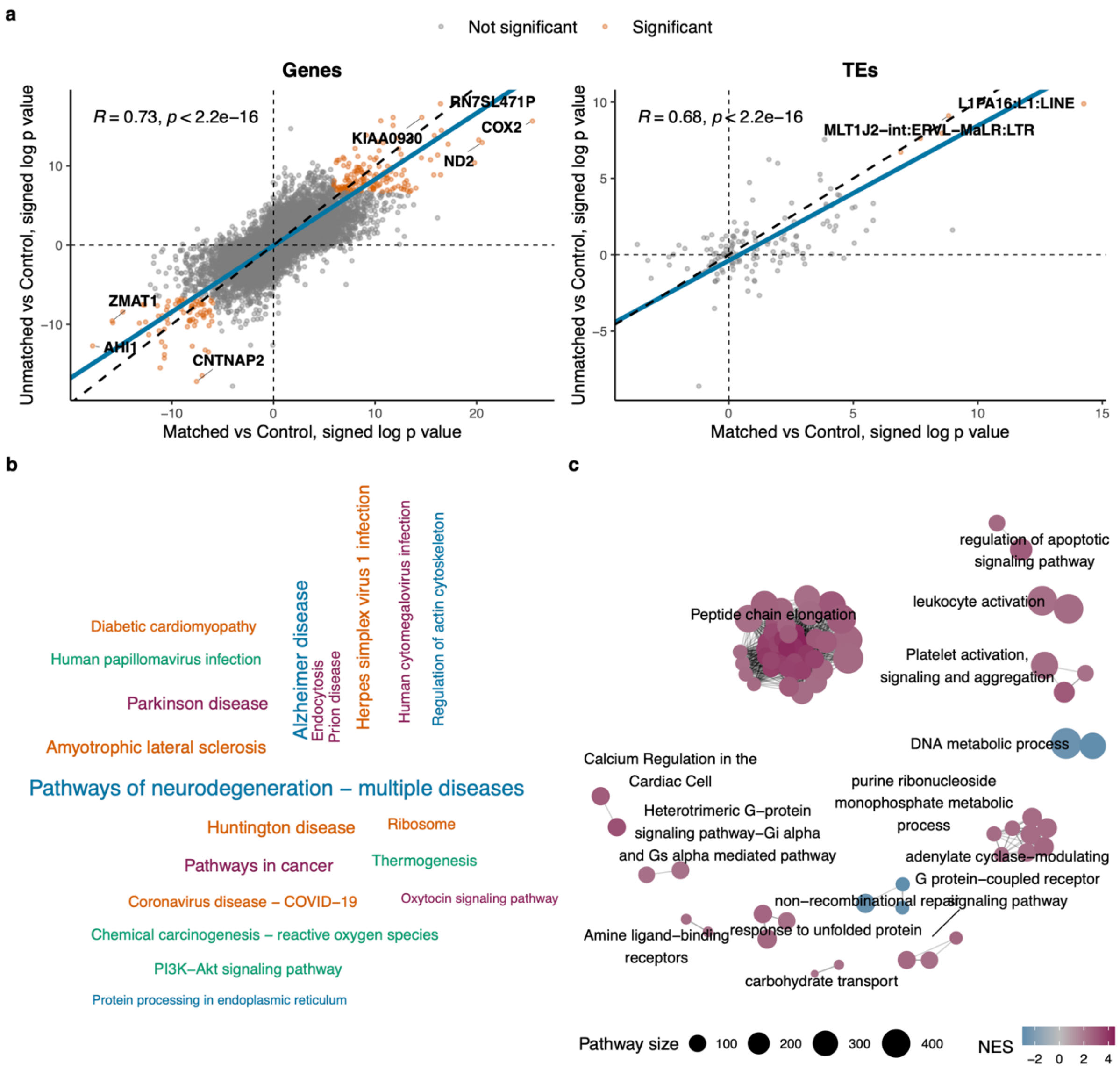


**Figure S2: Differential expression of genes and transposable elements (TEs) in sorted prefrontal cortex neurons comparing Unmatched PD to control brain samples.** (a) Correlation of gene expression and TE changes in Unmatched/Control and Matched/Control comparisons. Significant differentially expressed genes and TEs in both comparisons are indicated in orange. Labeled genes at the perimeter indicate the strongest effect in both datasets. Blue line was plotted using deming regression coefficients. (b) A word cloud of significantly enriched KEGG pathways obtained through GSEA of Unmatched/Control comparison. (c) Network and clustering of the significantly enriched pathways obtained through GSEA of Unmatched/Control comparison. GSEA, gene set enrichment analysis.

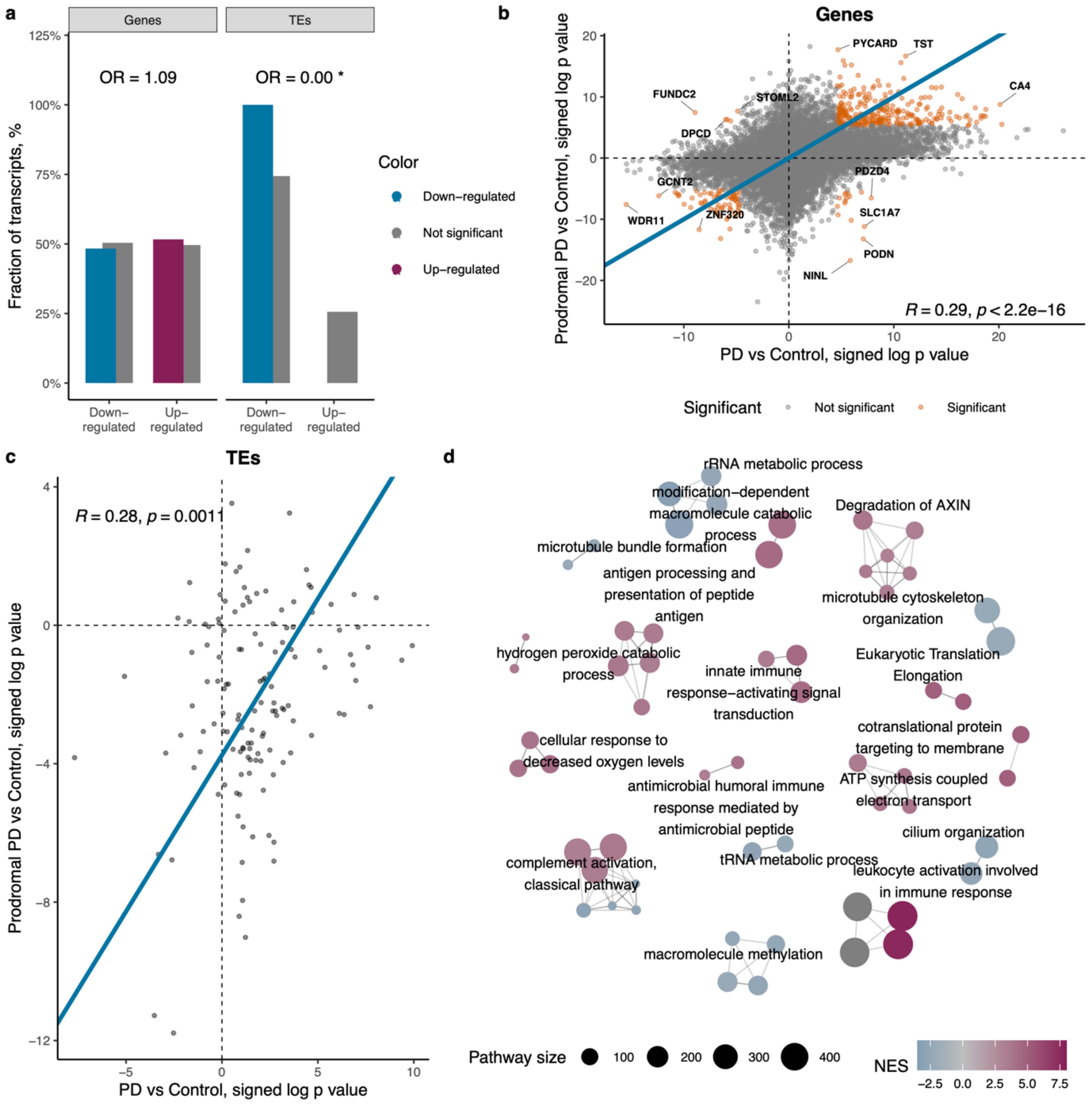


**Figure S3. Differential expression of genes and transposable elements (TEs) in the blood of prodromal PD patients. (a)** Fraction of up-regulated and down-regulated significant genes and TEs (colored fill) compared to the fraction of non-significant up-regulated and down-regulated genes as well as TEs (gray fill). **(b)** Correlation of gene expression changes in prodromal PD and PD patients compared to controls. Significant differentially expressed genes in both comparisons are indicated in orange. Labeled genes at the perimeter indicate the strongest effect in both datasets. Blue line was plotted using deming regression coefficients. **(c)** Correlation of TE expression changes in prodromal PD and PD patients compared to controls. Blue line was plotted using deming regression coefficients. **(d)** Network and clustering of the top 50 most up-regulated and 50 most down-regulated pathways obtained through gene set enrichment analysis. Each cluster is named after the most prominent pathway decided by the PageRank algorithm.

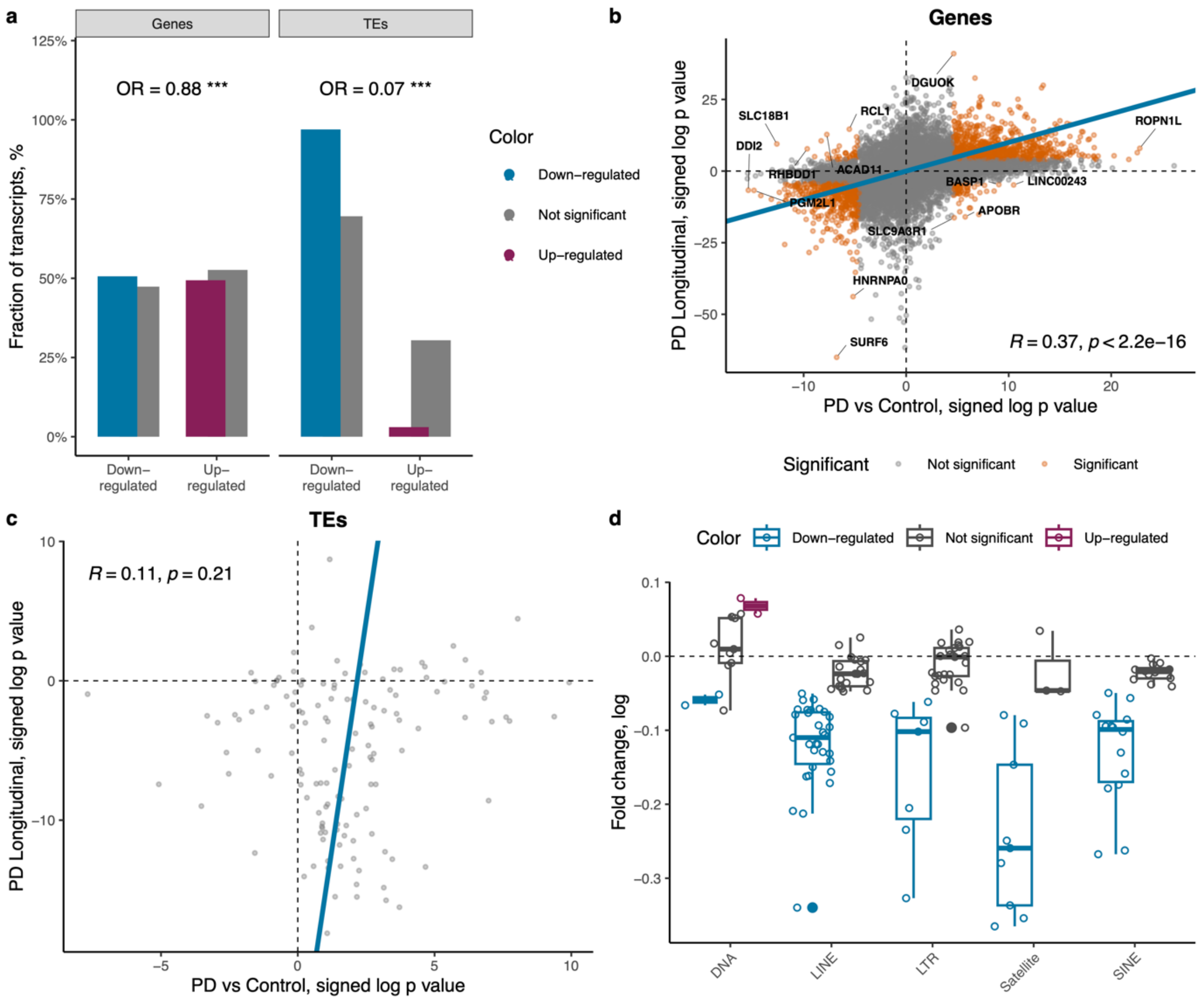


**Figure S4. Longitudinal expression changes of genes and TEs in blood of PD patients.** (a) Fraction of up-regulated and down-regulated significant genes and TEs (colored fill) compared to the fraction of non-significant up-regulated and down-regulated genes as well as TEs (gray fill). **(b)** Correlation of gene expression changes when comparing PD to Control samples and changes occurring longitudinally among the PD patients. Significant differentially expressed genes in both comparisons are indicated in orange. Labeled genes at the perimeter indicate the strongest effect in both datasets. Blue line was plotted using deming regression coefficients. **(c)** Correlation of TE expression changes when comparing PD to Control samples and changes occurring longitudinally among the PD patients. Blue line was plotted using deming regression coefficients. **(d)** Longitudinal change of TEs in PD stratified by class and colored by statistical significance.

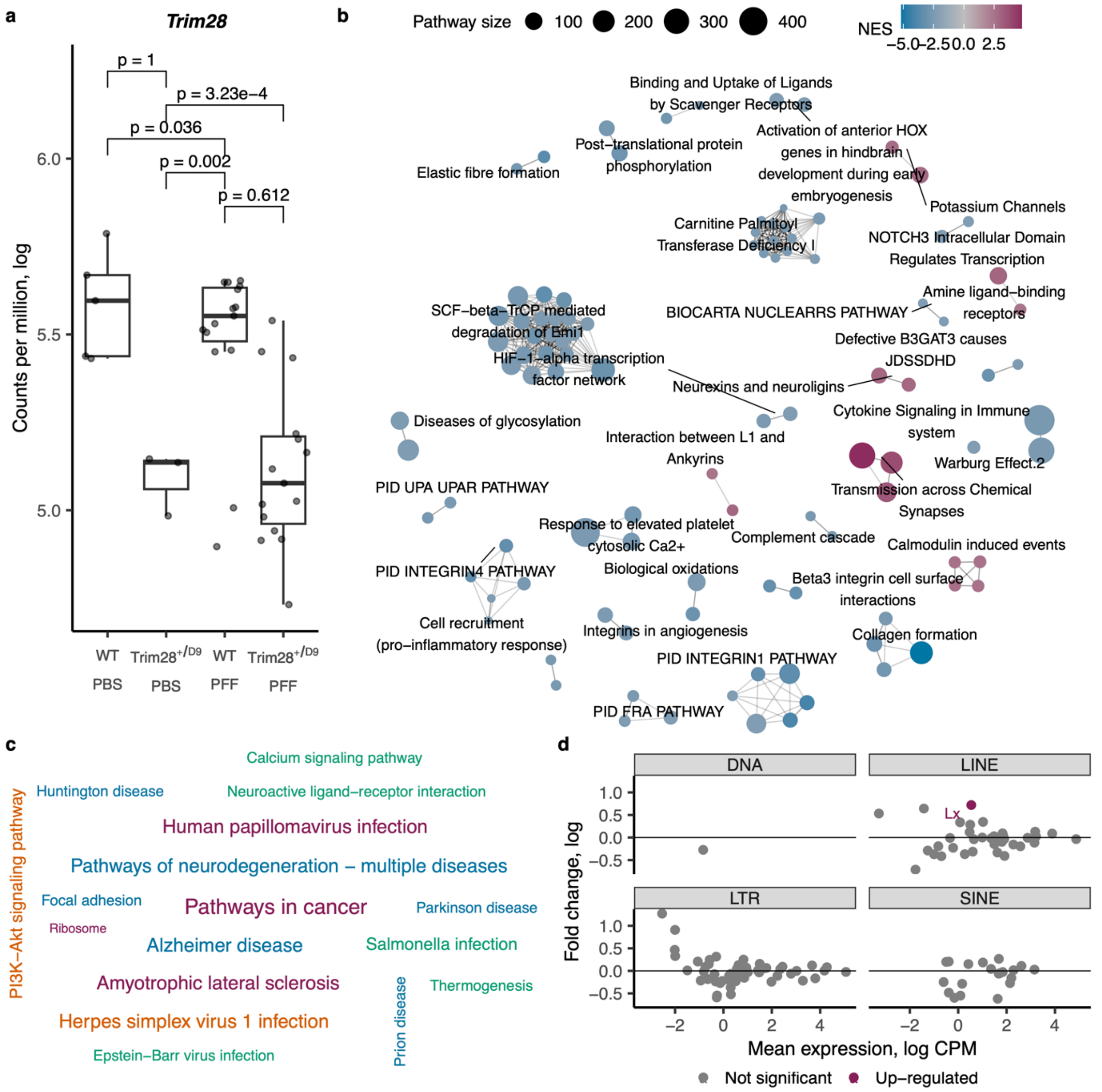


**Figure S5. Gene set enrichment analysis of Trim28 heterozygous knockout mouse samples.** **(a)** Trim28 down-regulation observed in Trim28 heterozygous mice. **(b)** Network and clustering of the significantly enriched pathways obtained through GSEA of Trim28$^{+/D9}$ to wild-type comparison. **(c)** Wordcloud of KEGG pathways obtained through GSEA of Trim28$^{+/D9}$ to wild-type comparison. **(d)** Mean expression and fold change of TEs stratified by class and colored by statistical significance in PFF injected mouse brain data.
